# Dynamics of *Mycobacterium tuberculosis* Ag85B revealed by sensitive ELISA

**DOI:** 10.1101/574996

**Authors:** Joel D. Ernst, Amber Cornelius, Miriam Bolz

## Abstract

Secretion of specific proteins contributes to pathogenesis and immune responses in tuberculosis and other bacterial infections, yet the kinetics of protein secretion and fate of secreted proteins in vivo are poorly understood. We generated new monoclonal antibodies that recognize the *M. tuberculosis* secreted protein, Ag85B, and used them to establish and characterize a sensitive ELISA to quantitate Ag85B in samples generated in vitro and in vivo. We found that nutritional or culture conditions had little impact on secretion of Ag85B, and that there is considerable variation in Ag85B secretion by distinct strains in the *M. tuberculosis* complex: compared with the commonly-used H37Rv strain (Lineage 4), *M. africanum* (Lineage 6) secretes less, and two strains from Lineage 2 secrete more Ag85B. We also used the ELISA to determine that the rate of secretion of Ag85B is 10-to 100-fold lower than that of proteins secreted by gram-negative and gram-positive bacteria, respectively. ELISA quantitation of Ag85B in lung homogenates of *M. tuberculosis* H37Rv-infected mice revealed that although Ag85B accumulates in the lungs as the bacterial population expands, the amount of Ag85B per bacterium decreases nearly 10,000-fold at later stages of infection, coincident with development of T cell responses and arrest of bacterial population growth. These results indicate that bacterial protein secretion in vivo is dynamic and regulated, and quantitation of secreted bacterial proteins can contribute to understanding pathogenesis and immunity in tuberculosis and other infections.

**Importance:** Bacterial protein secretion contributes to host-pathogen interactions, yet the process and consequences of bacterial protein secretion during infection are poorly understood. We developed a sensitive ELISA to quantitate a protein (termed Ag85B) secreted by *M. tuberculosis* and used it to find that Ag85B secretion occurs with slower kinetics than for proteins secreted by gram positive and gram negative bacteria, and that accumulation of Ag85B in the lungs is markedly regulated as a function of the bacterial population density. Our results demonstrate that quantitation of bacterial proteins during infection can reveal novel insights into host-pathogen interactions.

## Introduction

*Mycobacterium tuberculosis* employs secretion of specific proteins (estimated to include up to ∼25% of the bacterial proteome (1)) to survive, interact with host targets during infection (2, 3), manipulate its intracellular niche (2-2), and induce protective and pathogenic immune responses (8). Among the proteins that are most abundant in *M. tuberculosis* culture supernatants are members of a family of three closely related proteins, the antigen 85 (Ag85) complex consisting of Ag85A, Ag85B, and Ag85C (9). All three of these proteins exhibit enzymatic activity as mycolyl transferases, in which they catalyze transesterification reactions to synthesize trehalose monomycolate (TMM), trehalose dimycolate (TDM), and mycolated arabinogalactan (10, 11). Because of these enzymatic activities and their importance in constructing the mycobacterial envelope, Ag85A, Ag85B, and Ag85C have been considered potential drug targets for treatment of tuberculosis (10).

Due to their ability to induce adaptive CD4 and CD8 T lymphocyte responses in a broad range of vertebrate hosts, Ag85A and Ag85B have been investigated as antigens for tuberculosis vaccines, and are prominent components of at least seven candidate vaccines in various stages of development (http://www.aeras.org). Quantitative assays of mRNA have revealed that the genes encoding Ag85A and Ag85B (*fbpA* and *fbpB*, respectively) are expressed at high levels by bacteria in the lungs early after aerosol infection of mice, but their mRNA expression decreases markedly after the recruitment of antigen-specific effector T cells to the lungs (12-12). Consistent with the results of bacterial RNA quantitation, CD4 T cells specific for Ag85B are activated in the lungs between two and three weeks after infection of mice, but their activation markedly decreases concurrent with decreased bacterial expression of the *fbpB* gene (12, 15).

Despite considerable knowledge of the properties of the *fbpA* and *fbpB* genes and the antigenicity of their products, there is less information on the secretion, in vivo expression, and trafficking of the Ag85A or Ag85B proteins. Because of interest in Ag85B as a vaccine and/or diagnostic antigen, we generated new monoclonal antibodies to Ag85B and used them to establish a highly sensitive and specific ELISA. We then employed the ELISA in studies of secretion and trafficking of the Ag85B protein in vitro and in vivo.

## Results

### Generation and characterization of monoclonal antibodies to Ag85B

Monoclonal antibodies were generated using standard methods (16), from mice immunized with purified recombinant *M. tuberculosis* Ag85B (rAg85B), expressed in *E. coli*. Three monoclonal antibodies (mAbs), termed 710, 711, and 712, were selected for characterization. When examined by direct ELISA using wells coated with purified rAg85A, rAg85B, or rAg85C, and varying concentrations of antibody, all three mAbs recognized Ag85B, and yielded equivalent signals (Figure 1A). The three mAbs also recognized Ag85A in the direct ELISA, although recognition of Ag85A required higher antibody concentrations and reached lower maximum signal intensity than with Ag85B. mAb 710 generated a higher signal intensity, and at lower antibody concentrations compared with mAbs 711 and 712; the latter two mAbs also exhibited detectable binding to Ag85A when used at concentrations ≥ 1 µg/ml (Figure 1B). In contrast, mAbs 711 and 712 did not bind to Ag85C in direct ELISA at any antibody concentration examined, while mAb 710 bound Ag85C at antibody concentrations as low as 0.01 µg/ml (Figure 1C). Testing of the mAbs at a fixed concentration of 1 µg/ml on a dilution series of recombinant protein in direct ELISA, revealed that mAb 710 bound to Ag85A, Ag85B, and Ag85C at lower antigen concentrations than did mAb 711 or 712; mAbs 711 and 712 were indistinguishable in this assay. All three mAbs bound to Ag85B at lower antigen concentrations than required for binding to Ag85A or Ag85C (Figure 1D-E). Together, these results indicate that mAbs 710, 711, and 712 preferentially recognize Ag85B, although each of the mAbs also binds Ag85A and Ab85C when these antigens are present at high concentrations.

**Figure 1.**
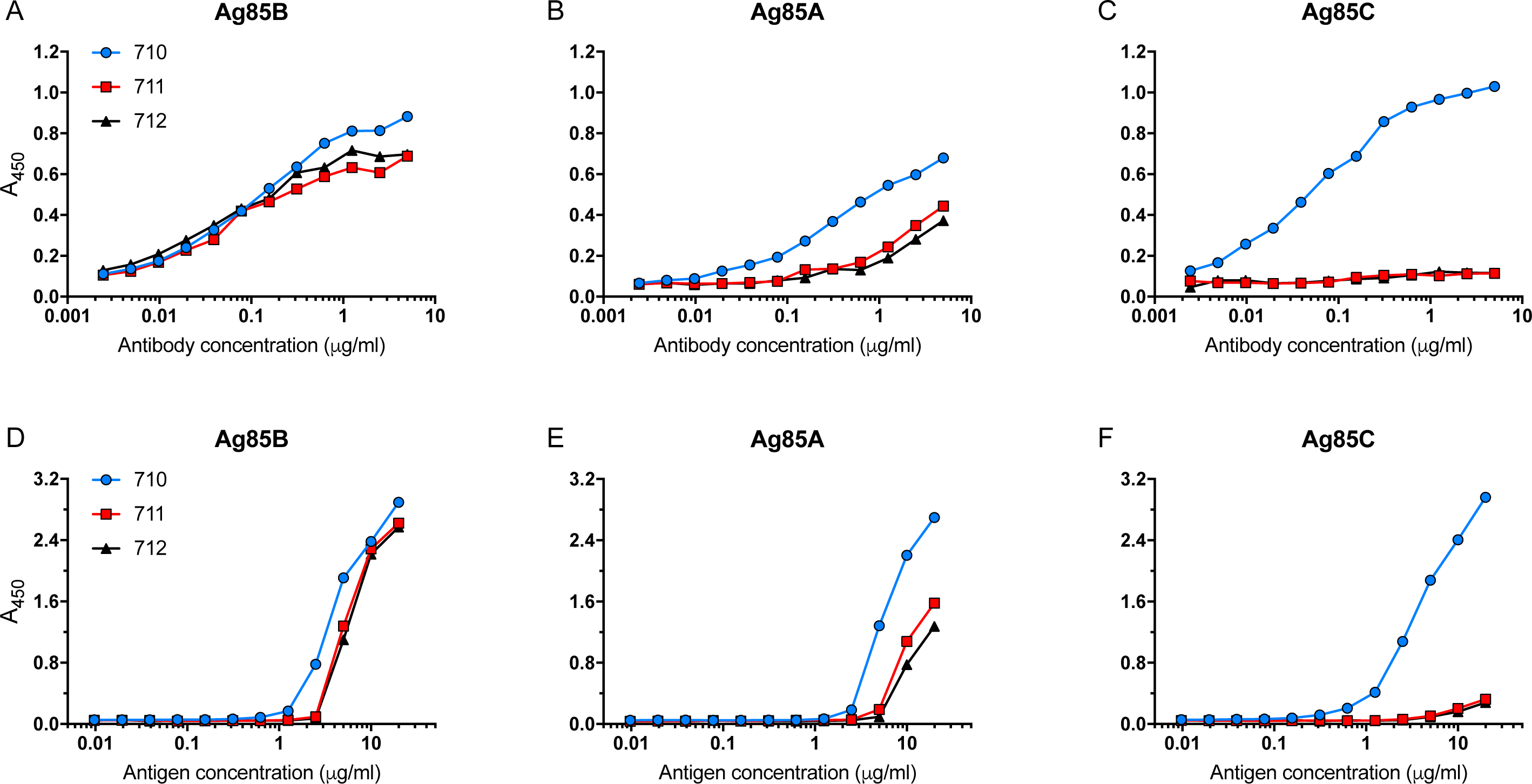
Recognition of Ag85A, Ag85B, and Ag85C by monoclonal antibodies 710, 711, and 712. Panels A-C: Wells of 96-well plates were coated with Ag85B (panel A), Ag85A (panel B), or Ag85C (panel C), each at 0.5 µg/ml. Monoclonal Abs 710, 711, or 712 were added at the indicated concentrations, and binding was detected after washing, using horseradish peroxidase (HRP)-conjugated goat anti-mouse IgG and TMB substrate. Data shown in Panels A-C are representative of three independent experiments with one technical replicate per experiment per condition. **Panels D-F:** Individual wells were coated with a dilution series of Ag85B (panel D), Ag85A (panel E), or Ag85C (panel F), starting at 20 µg/ml. Monoclonal Abs 710, 711, or 712 were added at a concentration of 1 µg/ml, and binding was detected after washing, using horseradish peroxidase (HRP)-conjugated goat anti-mouse IgG and TMB substrate. Data in panels D-F are representative of two independent experiments with one technical replicate per experiment per condition.

### Sandwich ELISA for quantitation of Ag85B

When the three mAbs were used in pairwise combinations, using one mAb for capture, and another mAb conjugated to horseradish peroxidase (HRP) for detection of bound antigen, the highest sensitivity for detection of Ag85B was obtained when mAb 710 was used as the capture antibody (Figure 2A). Although the differences were slight, sensitivity appeared to be greater when mAb 711 rather than mAb 712 was used as the detecting antibody. Compared with Ag85B capture by mAb 710, mAb 711 capture allowed detection of bound Ag85B by mAb 710 but not by mAb 712. When mAb 712 was used for capture, mAb 710 was able to bind Ag85B, but mAb 711 was not. Together, these data indicate that the epitopes recognized by mAbs 711 and 712 overlap or may even be identical, while the epitope recognized by mAb710 is distinct from those of mAbs 711 and 712. Despite detectable binding of all three mAbs to Ag85A in the direct ELISA (Figure 1B), none of the combinations of capture or detecting antibodies yielded a signal when Ag85A was used as the antigen in the sandwich ELISA (Figure 2B). Likewise, despite binding of mAb 710 to Ag85C by direct ELISA, none of the capture or detecting antibody combinations resulted in a detectable signal in the sandwich ELISA when Ag85C was used as the antigen (Figure 2C). Together, these results suggest that the epitope recognized by mAb 710 is at least partially shared by Ag85A, Ag85B, and Ag85C, while the epitope(s) recognized by mAbs 711 and 712 is more specific to Ag85B.

**Figure 2.**
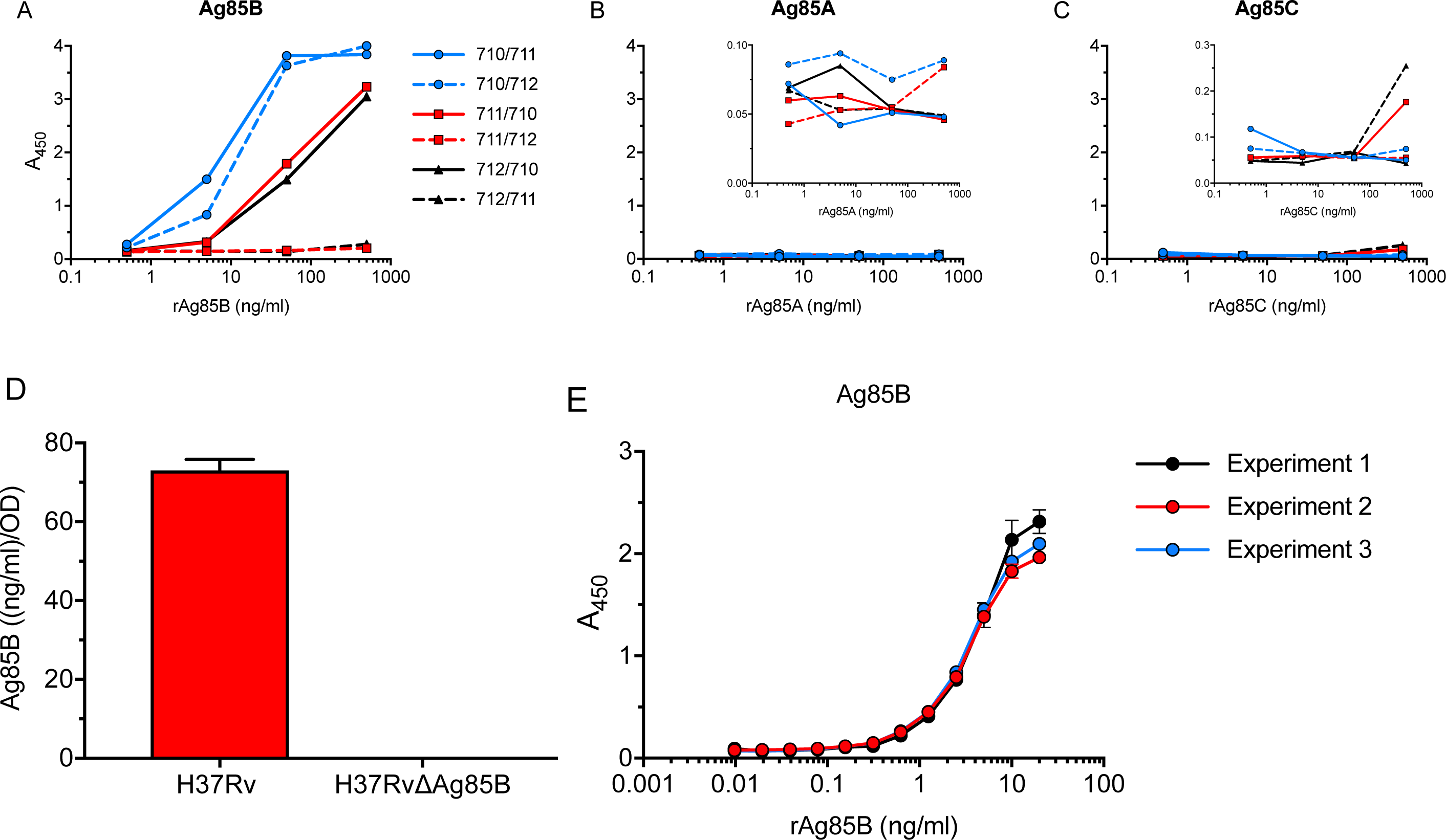
Characterization of sandwich ELISA for Ag85B. Each mAb was tested as the capture antibody, with either of the remaining two mAbs (conjugated to HRP) used for detection. A) Detection of rAg85B added to mAb-coated plates at the concentrations indicated on the X axis. B) Detection of rAg85A under the same conditions as in panel A. The inset shows the same data on a contracted scale, to reveal small differences. C) Detection of rAg85C under the same conditions as in panel A. The inset shows the same data on a contracted scale, to reveal small differences. D) Specificity of ELISA using mAb 710 for capture and HRP-conjugated mAb 711 for detection of antigen in culture filtrates of wild-type *M. tuberculosis* H37Rv or *M. tuberculosis* H37Rv with a targeted deletion of the gene encoding Ag85B. The H37Rv culture filtrate was diluted to give a signal in the linear range of the ELISA standard curve, while the H37RvΔAg85B culture filtrate was assayed undiluted. The date shown are mean (bar) and standard deviation (error bar) of biological triplicate values. E) Standard curves of rAg85B in Ag85B ELISA. Shown are curves of three independent experiments with two independent standard curves each. Shown are mean (dot) and standard deviation (error bar) when large enough to depict. Data shown in Panels A-C are representative of three independent experiments with one technical replicate per experiment per condition.

To confirm specificity of the ELISA with the combination of mAbs 710 and 711 using native *M. tuberculosis* proteins, we examined culture filtrates of wild-type H37Rv and of our previously-characterized Ag85B-deficient (*fbpB* null) mutant strain of H37Rv (12). This yielded no detectable signal in the ELISA when culture filtrates of the Ag85B-deficient bacteria were examined, despite the detection of an abundant signal when wild-type culture filtrates were examined (Figure 2D). Since the Ag85B-null bacteria retain the capacity to synthesize and secrete Ag85A and Ag85C (17), these results provide further evidence that the sandwich ELISA with mAbs 710 and 711 is highly specific for Ag85B.

The assay in this form, with mAb 710 as the capture antibody and HRP-coupled mAb 711 as the detection antibody, has been run in our laboratory >30 times to determine Ag85B content in various samples. Each assay comprised at least one standard curve with rAg85B per ELISA plate, starting at 10 ng/ml or above and including 2-fold dilutions to loss of signal (generally,≤20 pg/ml). Three representative assay standard curves are shown in Figure 2E. Ag85B in samples was quantitated based on either a dose-response sigmoidal regression curve or a linear regression in the linear proportion of the curve (0 ng/ml – 2.5 ng/ml). With both methods regression curves had R^2^ values > 0.99. When necessary, samples were diluted for ELISA to be in linear range of the standard curve and either at least two technical replicates or biological replicates (when available) were run.

### Carbon source effects, strain-dependent variation, and kinetics of Ag85B secretion

We used the ELISA to examine whether Ag85B secretion is governed by bacterial growth conditions. When grown in rich broth (Middlebrook 7H9 medium with 10% albumin-dextrose-catalase), a 10-fold difference in Tween 80 concentration (0.5% vs 0.05%) did not affect the amount of Ag85B secreted by *M. tuberculosis* H37Rv in over a 24 h period (Figure 3A). When H37Rv was grown in minimal defined media (Sauton’s medium, with 0.05% Tween 80, 0.5% BSA with 0.05% tyloxapol), or 7H9 broth with the addition of 0.2% acetate, dextrose or glycerol as carbon sources, the addition of acetate or glycerol did not alter Ag85B secretion, whereas 0.2% dextrose increased Ag85B secretion approximately 2-fold (Figure 3A). These results indicate that bacterial metabolism as dictated by alternative carbon sources can affect the rate and/or amount of Ag85B secretion, but that the concentration of Tween 80 had little measurable effect.

**Figure 3.**
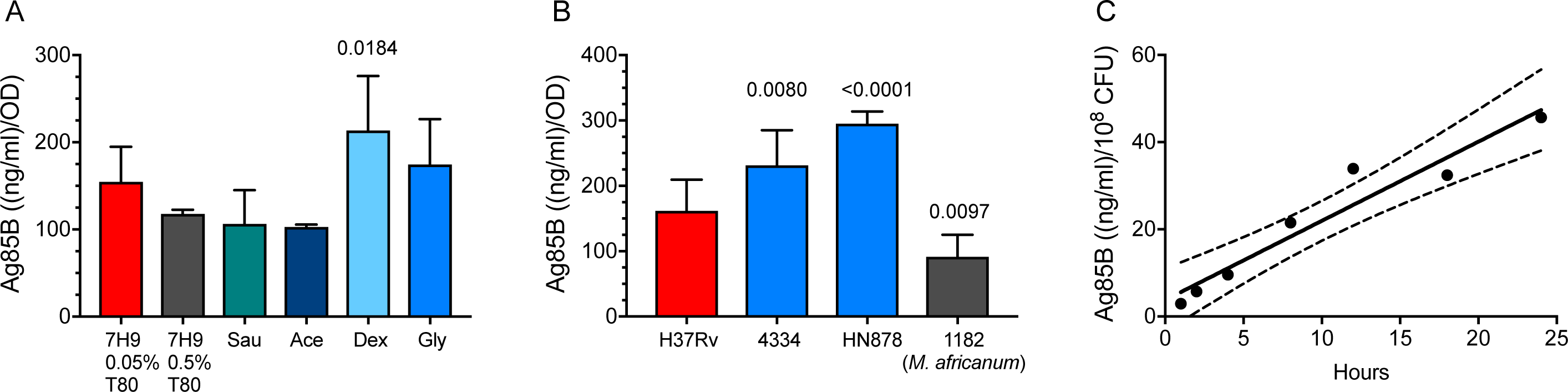
Effects of carbon source and *M. tuberculosis* strain, and kinetics of Ag85B secretion. A) Effects of media composition and carbon source on Ag85B secretion. Mid-log phase bacteria (A600 = 1.0) grown in 7H9 medium were washed 5 times, then diluted 10-fold into the indicated media. After 24 h, the A600 was determined for each culture, and sterile filtrates were prepared for analysis by mAb 710-711 ELISA. Results of Ag85B quantitation by ELISA were normalized by the A600 of the individual culture. 7H9 0.05% T80: Middlebrook 7H9 with 10% ADC and 0.05% (v/v) Tween 80; 7H9 0.5% T80: same as preceding, but with 0.5% Tween 80; Sau: Sauton’s medium with 0.05% Tween 80; Ace: 7H9 media with acetate; Dex: 7H9 media with dextrose; Gly: 7H9 media with glycerol. Show are mean (bar) and standard deviation (error bar) of biological triplicate values. Statistical comparison was done using one-way ANOVA; the adjusted p value after Dunnett’s post-test for multiple comparisons was applied is shown for the effect of dextrose. Other effects were not significant after adjusting for multiple comparisons. B) Mycobacterial strain-dependent variation of Ag85B secretion. Mid-log phase cultures were collected and the bacteria from each strain were washed and resuspended in fresh 7H9 broth. After 24h, the bacteria were pelleted by centrifugation and Ag85B in culture filtrates was quantitated by ELISA. Show are mean (bar) and standard deviation (error bar) of biological triplicate values. The adjusted p values shown are for comparison of each strain with H37Rv, and were determined by one-way ANOVA with Dunnett’s post-test. C) Kinetics of Ag85B secretion by *M. tuberculosis* H37Rv. Mid-log phase growing H37Rv was washed, added to fresh 7H9 medium at 1 x 10^8^ CFU/ml, and incubated for 24 h. Individual cultures were sampled at the designated time points and culture filtrates were assayed by ELISA. Statistical analysis by linear regression was determined using Prism 7. Dashed lines indicate 95% confidence interval. Figure A shows one representative of three independent experiments. Figures B and C show data of experiments each done once.

We previously reported that *M. africanum* expresses lower levels of Ag85B than *M. tuberculosis* H37Rv, as detected by immunoblotting and by the magnitude of in vivo antigen-specific CD4 T cell responses (18). To verify this result and determine whether other strain-dependent variation in Ag85B secretion exists, we used the Ag85B-specific ELISA to assay culture filtrates from phylogenetically distinct bacterial isolates. This confirmed that *M. africanum* secretes significantly lower quantities of Ag85B compared with *M. tuberculosis* H37Rv (Figure 3B). We also found that two distinct isolates of *M. tuberculosis* from Lineage 2 (includes the Beijing family) secrete significantly higher quantities of Ag85B than H37Rv (Figure 3B). These results indicate substantial variation in secretion of Ag85B, according to the bacterial strain, possibly in a lineage-dependent manner.

To further characterize the properties of Ag85B, we determined the kinetics of its secretion by *M. tuberculosis* H37Rv. Washed mid-log phase bacteria were suspended in fresh 7H9 media at 10^8^ CFU/ml, and culture filtrates were harvested at multiple intervals and assayed by ELISA. The rate of accumulation of Ag85B in culture filtrates was linear (r^2^ = 0.93) during the 24-hour period of sampling, at 1.8 ± 0.2 ng/ml/hr (Figure 3C). Since the mature Ag85B protein has a molecular weight of 34,580, this is equivalent to 52 fmol per 10^8^ CFU/hr, or approximately 300 molecules secreted per bacterial cell/hr.

### Cell-free Ag85B during infection in vivo and in vitro

Despite their biological activity and roles in pathogenesis and immune responses, little is known of the in vivo fate or distribution of secreted *M. tuberculosis* proteins, especially during infection. Therefore, we examined supernatants of lung homogenates obtained at various intervals after infecting C57BL/6 mice with *M. tuberculosis* H37Rv by aerosol. We found that Ag85B was detectable in lung homogenate supernatants in some, but not all, mice as early as 4-8 days post-infection, followed by a progressive increase between 14 and 21 days post-infection (Figure 4A). Since the samples for assay were taken during the progressive growth phase of the bacteria in the lungs, the quantity of Ag85B detected is a function of the amount secreted by each bacterium and by the number of bacteria, which increases progressively until approximately 21 days post-infection (19). Therefore, we normalized the concentration of Ag85B in lung homogenates by the number of bacteria present in the lungs at each time point sampled. This revealed a progressive decrease in the amount of Ag85B relative to the bacterial population size, commencing between 8 and 11 days post infection (Figure 4B). This result is consistent with results of Ag85B RNA quantitation on bacterial populations in the lungs of immunocompetent mice (12-14), and with results of studies indicating that activation of Ag85B-specific CD4 T cells in vivo decreases markedly after 2-3 weeks of infection, due to limited availability of antigen (12, 15).

**Figure 4.**
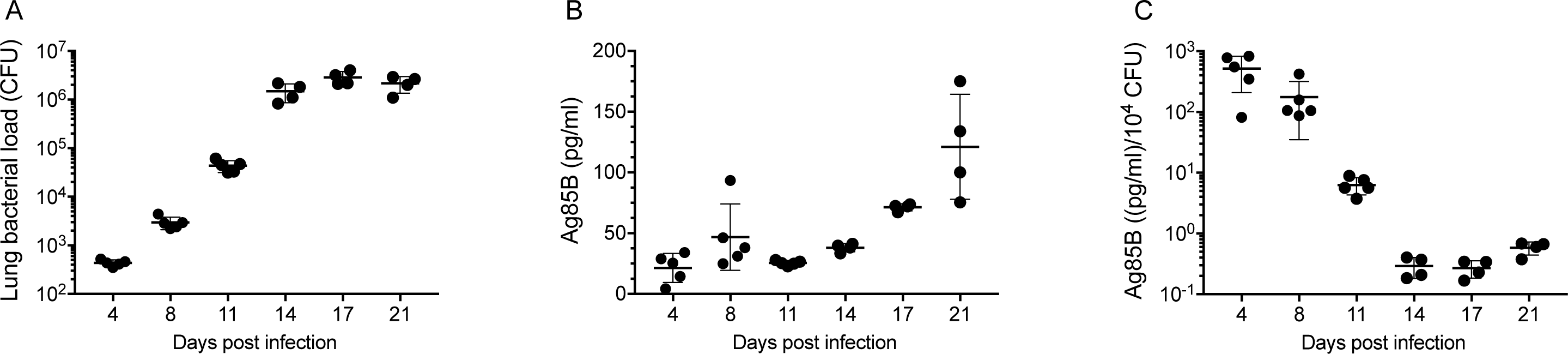
Quantitation of Ag85B in supernatants of lung homogenates from mice infected with *M. tuberculosis* H37Rv. A) *M. tuberculosis* colony-forming units (CFU) in the lung samples used for assays in panel B. B) Ag85B concentrations quantitated by ELISA in lung homogenate supernatants; C) Ag85B concentrations as in Panel B, normalized by the number of bacteria (CFU) in the same lung homogenates. Shown are individual data points, mean ± standard deviation. This experiment was done once.

There are several potential sources of free Ag85B in the lungs. First is that some of the bacteria may be extracellular and may secrete Ag85B directly to the intercellular spaces in lung tissues. Second is that, as we have recently reported, Ag85B and other secreted mycobacterial proteins can be exported from infected cells by a vesicular transport pathway (20, 21). A third potential mechanism is release of Ag85B associated with bacterial membrane vesicles (22) and/or exosomes shed by infected cells (23, 24). A fourth possible mechanism is that Ag85B synthesized by intracellular bacteria may be released from dying infected cells. We investigated the latter possibility, using bone marrow-derived dendritic cells infected with *M. tuberculosis* H37Rv. Under conditions of the multiplicities of infection, bacterial strain, and time points used, we observed a range of loss of cell viability as reflected by luminescence assay of ATP in cell lysates, after harvesting conditioned medium for assay of Ag85B by ELISA. This revealed a correlation between loss of cell viability and the quantity of Ag85B in conditioned media (r^2^ = 0.5777; p < 0.0001) (Figure 5), indicating that Ag85B can be released from dead and/or dying infected cells in a form that remains detectable by ELISA. This suggests that death of infected cells can be a source of the Ag85B detected in cell-free homogenates of the lungs of infected mice, as in Figure 4.

**Figure 5.**
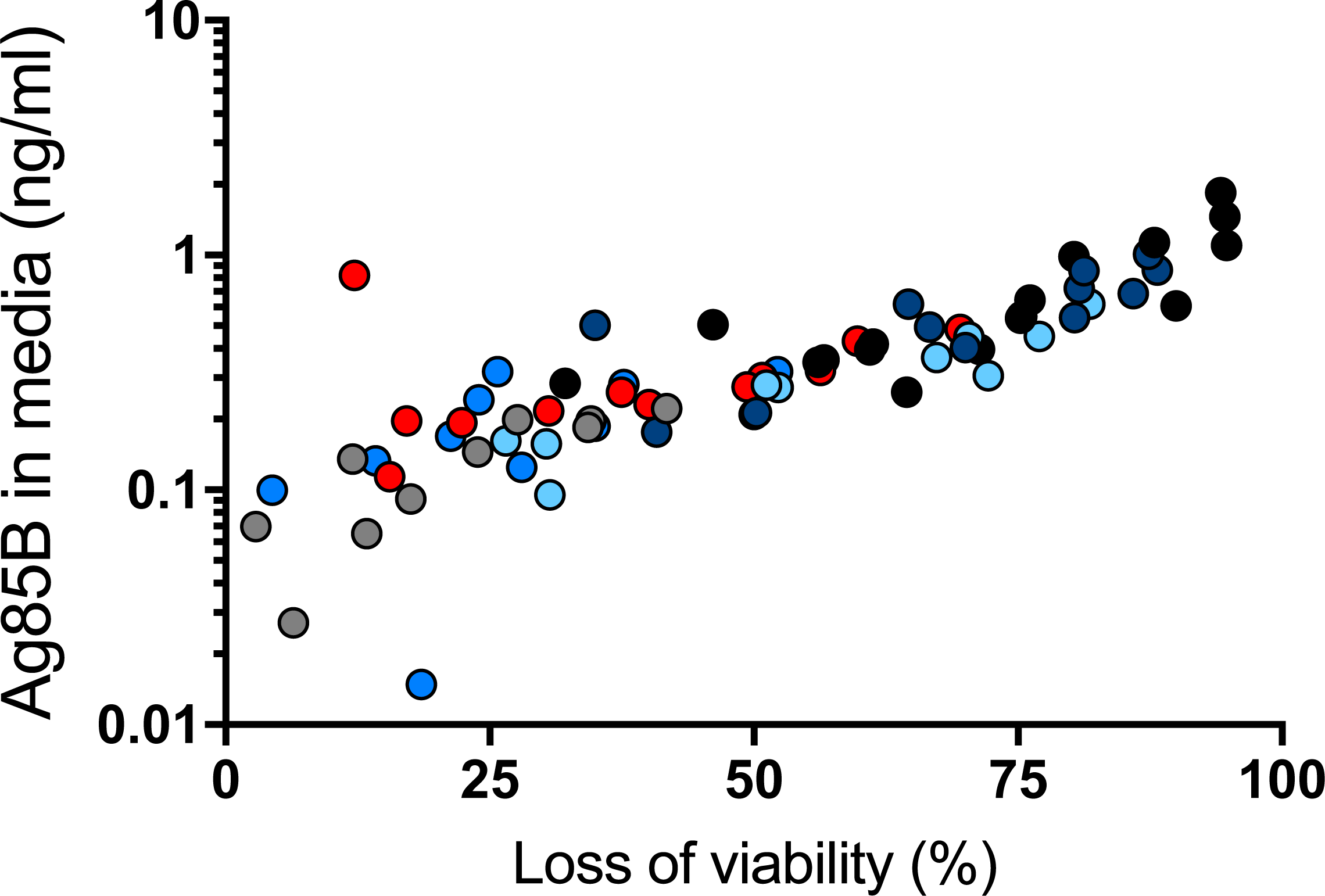
Death of *M. tuberculosis*-infected primary dendritic cells releases Ag85B. Bone marrow-derived dendritic cells were infected with *M. tuberculosis* H37Rv at different multiplicities of infection (1, 2, 4 and 8). At designated time points (12 h grey dots, 16 h medium blue dots, 24 h red dots, 36 h light blue dots, 48h dark blue dots), medium was removed for quantitation by ELISA, and dendritic cell viability was assessed by luminescence assay of ATP of cell lysates. Results of the two assays on a given sample were plotted as a data point, and Pearson correlation was determined for the whole dataset. Shown are representative results of two similar experiments.

## Discussion

In this work, we developed three new murine monoclonal antibodies by immunizing mice with recombinant *M. tuberculosis* Ag85B, and used them to develop a sensitive and specific ELISA for quantitation of Ag85B in various biological samples. We found that, compared with the H37Rv strain from the *M. tuberculosis* complex lineage 4, two isolates from lineage 2 secreted greater quantities, while a lineage 6 (*M. africanum*) isolate secreted lesser quantities of Ag85B. Since Ag85B plays a role in synthesis of trehalose dimycolate, and trehalose dimycolate is a potent proinflammatory mediator during *M. tuberculosis* infection (25-28), the results suggest that differences in Ag85B secretion may contribute to strain-dependent differences in the proinflammatory properties of distinct strains and lineages in the *M. tuberculosis* complex (29). Strain variability in secretion may also influence the frequency and magnitude of T cell responses to Ag85B, which may provide an explanation for the finding that fewer human subjects exhibit detectable responses to Ag85B compared with other antigens such as ESAT-6 and/or CFP-10 (30, 31). Likewise, bacterial strain-dependent variation in secretion of Ag85B may influence the protective efficacy of TB subunit vaccines that include Ag85B: even if a vaccine generates immune responses to Ag85B, vaccination may provide little protection against infection with mycobacterial strains that express and secrete lesser quantities of Ag85B. In studies of subunit vaccines that include Ag85B, it may be informative to characterize Ag85B expression and secretion in the isolates of *M. tuberculosis* from subjects that develop breakthrough infections despite apparently-appropriate immune responses to the vaccine antigen.

We also used the ELISA to determine the rate of secretion by *M. tuberculosis* H37Rv in broth culture as approximately 300 molecules of Ag85B per bacterial cell per hour. This is 10 to 100-fold lower than the rate of translocation of *E. coli* proOmpA (4.5 mol/min) (32), or the secretion of Staphylococcal toxic shock toxin (1.1 x 10^4^ molecules/CFU/hr) (33). To fully understand the basis for this difference will require determination of the rates of protein synthesis in *M. tuberculosis* compared with that of other bacteria, and characterization of the factors and mechanisms that determine the rate of protein secretion by distinct secretion systems in bacteria. Given the important roles of secreted protein virulence factors in *M. tuberculosis* and other bacterial pathogens, better understanding of protein secretion may reveal new targets for therapeutic modulation and reduction of disease and pathogen transmission.

A third major finding in these studies is that *M. tuberculosis* Ag85B can be found in homogenized lung tissue supernatants from infected mice. This finding has several potential implications. If data from studies in human subjects provide supportive data, then detecting Ag85B in respiratory secretions may provide a rapid and economical approach to diagnosis of pulmonary tuberculosis and may also be useful in monitoring responses to treatment. Another is that the presence of extracellular Ag85B in lung tissue may make Ag85B available for uptake and processing and presentation by uninfected dendritic cells and macrophages in the lungs. Since we have reported that direct recognition of infected cells is required for optimal CD4 T cell control of intracellular *M. tuberculosis* (34), acquisition and presentation of Ag85B by uninfected cells in the lungs may provide antigen-loaded decoys for Ag85B-specific CD4 T cells that reduce the frequency of recognition of infected cells by those T cells. This is consistent with our recent finding in mouse lungs (35) and the finding in lungs of *M. tuberculosis-*infected rhesus macaques (36) that only a small fraction of the T cells in the lungs are in close contact with *M. tuberculosis*-infected cells. Our associated finding in this work that death of *M. tuberculosis*-infected host cells can be associated with release of Ag85B to the extracellular space indicates that the presence of Ag85B (and other mycobacterial proteins) in extracellular tissue compartments in the lungs may be the consequence of secretion by extracellular bacteria, vesicular transport from infected cells (21), carriage by exosomes (37,38), and release by dying or dead host cells. Since Ag85B (39, 40), ESAT-6 (41) and other mycobacterial proteins have been reported to act on host cells to modulate inflammation, our findings provide additional evidence for the plausibility that extracellular mycobacterial proteins contribute to the pathogenesis of TB, and may be susceptible to therapeutic modulation.

## Materials and Methods

### Ethics statement

All animal experiments were done in accordance with procedures approved by the New York University School of Medicine Institutional Animal Care and Use Committee (Laboratory Animal Care Protocol 150502-01), which conformed to the guidelines provided by the Guide for the care and Use of Laboratory Animals of the National Institutes of Health.

### Bacterial strains

The stocks of *M. tuberculosis* H37Rv, H37Rv:Δ*fbpB* (Ag85B-null), 4334, and *M. africanum* 1182 used in our laboratory and for these studies have been previously described (19, 42-44). The *M. tuberculosis* HN878 strain was obtained from BEI Resources.

### Recombinant Ag85B

RV1886c-ss.pET23b was transformed into *E. coli* BL21(DE3) pLysS (Invitrogen) and induced with 0.8 mM IPTG for 4 h at 37° C. Cultures were lysed with 50 mM Tris pH 7.0, 150 mM NaCl, lysozyme, benzonase, and 1 mM PMSF for 30 minutes at 22°C on an orbital shaker. The lysate was spun 10,000 x g for 30 minutes, sterile filtered, and loaded onto an AKTA FPLC His-Trap column (GE). The column was washed with 50 mM imidazole, and recombinant protein was eluted with 250 mM imidazole. Purity was assessed by SDS-PAGE.

### mAb generation and selection

Balb/C mice were immunized with purified recombinant Ag85B (100 µg/mouse x 2 injections followed by 50 µg/mouse for 2 additional injections) subcutaneously in TiterMax Gold (TiterMax, Norcross, GA, USA), followed by 50 µg/mouse given intravenously 3 days before harvesting and using spleen cells in fusions with P3X63Ag8 myeloma cells. Hybridoma supernatants were screened for recognition of rAg85B-coated wells by ELISA.

### Direct ELISA for characterization of individual mAbs

Ag85A (BEI Resources NR-14871), Ag85B (our purified recombinant), or Ag85C (BEI Resources NR-14858) were used to coat wells at 0.5 µg/well in PBS and incubated overnight at 4°C. The plates were washed 3 times with PBS pH 7.4 with 0.05 % Tween 20, and blocked with PBS containing 1.0 % BSA for 1 hour. A starting concentration of 5 µg/ml of mAb 710, 711, or 712 was serially diluted and 200 μl/well incubated at room temperature for 2 hours. Plates were washed 5 times and incubated with goat anti-mouse IgG HRP (MP Biomedicals) for 1 hour at room temperature. Plates were washed 7 times and developed with TMB substrate according to the manufacturer’s instructions (BD). The reaction was stopped with 2M sulfuric acid and absorbance read at 450 nm with a Synergy H1 micro plate reader (BioTek).

### Mycobacterial culture filtrates

*M. tuberculosis* strains H37RV, 4334, and HN878, or *M. africanum* strain 1182, were inoculated from a 1 ml frozen stock of approximately 3x10^8^ CFU/ml into 10 ml of Middlebrook 7H9 media supplemented with 0.5 % albumin, 0.2 % dextrose and 0.3 mg/100 ml catalase (in the text, termed 7H9) and grown to late log phase. The cultures were then passaged and grown once to mid log phase. Cultures were then collected and spun at 150 x g. The collected supernatant was then spun at 3,750 x g for 5 minutes and washed with PBS. The pellets were then resuspended in fresh 7H9 media and grown for 24 h. The cultures were pelleted and the media was sterile filtered.

### Sandwich ELISA development and optimization

Individual mAbs were used at 2.5 µg/ml in 50 µl 0.05 M carbonate-bicarbonate buffer to coat wells overnight at 4° C. The plates were washed 3 times with PBS (pH 7.4) with 0.5% Tween 20 and blocked with PBS containing 1.0% BSA for 1 hour. Antibodies for detection were conjugated to HRP according to manufacturer’s recommendations (Abcam). Briefly, 1 µl of Modifier was mixed with 10 µl of antibody and then added to a vial of HRP Mix. The vial was incubated overnight at room temperature in the dark. After incubation, 1 µl of Quencher was added and mixed. The conjugated antibodies were diluted to 0.5 µg/ml in PBS with 1% BSA and stored in aliquots at -20 °C until first use. Upon thawing, conjugated antibodies were stored at 4 °C.

For ELISA, samples were added to plates coated with the designated antibody and incubated at room temperature for 2 hours, then plates were washed 5 times. The labeled, detecting antibody was added and the plates were incubated at room temperature for 1 hour. Plates were washed 7 times and developed with substrate according to the manufacturers’ instructions (BD). The reaction was stopped with 2M sulfuric acid and absorbance read at 450 nm.

### Sandwich ELISA application

The routine sandwich ELISA used the procedures as described above, with mAb 710 as the capture antibody used to coat wells, and HRP-conjugated mAb 711 used to detect bound antigen. Signal generation for the HRP reaction used TMB substrate (Thermo Scientific).

### Effects of medium and carbon source

After washing in PBS, bacteria were resuspended in 6 ml of distilled water and split among 6 bottles. 10 ml of media was added to the bottles. The media were Middlebrook 7H9 with 10% ADC and either 0.05% or 0.5% Tween 80; or Sauton’s broth with 0.05% Tween 80, 0.5% BSA, and 0.05% tyloxapol, with 0.2% (w/v) acetate, glycerol, or dextrose (adapted from (45)), and bacteria were incubated for 24 h prior to preparation of culture filtrates.

### Preparation of lung homogenate supernatants from M. tuberculosis-infected mice

C57BL/6 mice were infected by the aerosol route with ∼60 CFU/mouse of *M. tuberculosis* H37Rv; lungs were harvested and single-cell suspensions were prepared as previously described (46). After sampling the cell suspensions for determination of bacterial CFU, an aliquot of each sample was sterile-filtered, and Ag85B concentration determined by ELISA. *Culture, infection, and analysis of bone marrow-derived dendritic cell death and release of Ag85B.*

Bone marrow-derived dendritic cells (BMDC), generated as previously described (20), were seeded in 96-well tissue culture treated plates (Corning) at 2x10^6^/well and rested for 2 hours, then infected overnight with different multiplicities of infection (1, 2, 4 and 8) with *M. tuberculosis* H37Rv, treated with Amikacin (200 μg/ml, 40min) in BMDC medium (RPMI1640 supplemented with 10% heat-inactivated FBS, 2 mM L-glutamine, 1 mM sodium pyruvate, 1xβ-mercaptoethanol, 10 mM HEPES and 12 ng/ml recombinant mouse GM-CSF), washed three times in PBS and further cultured in fresh BMDC medium. CM was harvested 16, 24, 34 and 48 hours later, sterile filtered and Ag85B in CM quantified by sandwich ELISA. At each harvest time point infected BMDC were assayed for cell death by CellTiter-Glo (Promega) according to the manufacturer’s instructions and signal read as Luminescence with a Synergy H1 micro plate reader (BioTek). For each harvest time point signal from uninfected cells was considered as 100 % viability for determination of loss of viability of infected cells.

### Statistical analyses

All statistical analyses were performed using Prism 7 (Graphpad). The specific tests used for data analysis are specified in the individual figure legends.

## Acknowledgements

We thank Smita Srivastava, Ph.D., for her contributions to the initial stages of this project.

